# Dexmedetomidine mediates microRNA-185 to suppress ovarian cancer growth via inhibiting the SOX9/Wnt/β-catenin signaling pathway

**DOI:** 10.1101/2020.04.15.042135

**Authors:** Hang Tian, Lei Hou, Yumei Xiong, Qiuju Cheng

## Abstract

**Objective:** Cervical cancer (CC) is a serious health problem, causing heavy burden each year worldwide. Long non-coding RNA (lncRNA) is implicated in CC progression. This study aims to clarify the underlying mechanism of lncRNA HAND2-AS1 in CC cellular process.

**Method:** LncRNA expression in CC tissues and adjacent tissues was analyzed via Gene Expression Omnibus microarray. The levels of HAND2-AS1, microRNA (miR)-21 and ALEX1 in 54 cases of CC tissues, adjacent tissues, CC cell lines and normal breast cells were detected. The malignant biological behaviors of CC cells were detected after overexpressing HAND2-AS1 or inhibiting miR-21 expression. Bioinformatics analysis, luciferase reporter assay and RNA-pull down assay were performed to verify the relationship among HAND2-AS1, miR-21 and ALEX1. The effects of HAND2-AS1 on CC cell growth *in vivo* were verified by nude mice transplantation experiment.

**Results:** HAND2-AS1 was lowly expressed in CC tissues and cells, and HAND2-AS1 functioned as a competitive endogenous RNA of miR-21 and enhanced ALEX1 expression and downregulated JAK1/STAT3 pathway to inhibit malignant biological behavior of CC cells. Overexpression of HAND2-AS1 or silencing of miR-21 can inhibit the malignant biological behavior of CC cells. Moreover, overexpression of HAND2-AS1 can inhibit the growth of CC cells *in vivo*.

**Conclusion:** Our data supported that lncRNA HAND2-AS1 promoted ALEX1 expression through competitively binding to miR-21, thereby inhibiting the proliferation and epithelial-mesenchymal transformation and promoting apoptosis of CC cells.

## Introduction

Ovarian cancer (OC) is a common deadly gynecological malignancy which accounts for about 3% of all cancer occurrences in female, with 240,000 cases newly found and 150,000 cases dead annually around the world (Hua and Li, 2018). Owing to the lack of early specific symptoms, OC is always diagnosed at advanced stages that hard to be cured by simple surgical resection, and frequent relapse occurs during the clinical course of OC (Bareiss and Paczulla, 2013), which in turn contributes to unfavorable prognosis and poor survival rate of OC patients ((Nikpayam and Tasharrofi, 2017; Ozga and Aghajanian, 2015). Despite the efficient surgical debulking and the application of different anticancer drug combinations, the overall 5-year survival rate of advanced OC patients was less than 40% with only modest improvement over decades (Gao and Liu, 2017; Varughese and Cocco, 2011). Investigating biomarkers enabling early diagnosis of OC and developing novel therapies continue to be fundamental goals for researchers in the field of OC.

Dexmedetomidine (DEX) is a strong α_2_-adrenoceptor agonist that is widely used for its anxiolytic, sedative, and analgesic attributes (Weerink and Struys, 2017). DEX could serve as an adjuvant analgesic in cancer patients undergoing surgical and nonsurgical procedures (Mohta and Kalra, 2016; Roberts and Wozencraft, 2011). Meanwhile, the positive effects of DEX such as reduction in respiratory depression and hypotension, decrease of lung and kidney damage and neuronal protection have been well-established (Endesfelder and Makki, 2017; Wang and Wu, 2018; Zhang and Jia, 2018). Considering these effects of DEX, one can presume beneficial roles of DEX in cancer patients. MicroRNAs (miRNAs) are small noncoding RNAs which regulate the translation of diverse genes and serve as ideal biomarkers for cancer prognosis (Ji and Shi, 2009). miRNAs play crucial roles in various essential biological processes including cell growth, differentiation, inflammation and tumor development, whose abnormal expression could lead to several diseases including cancer (Palma and Gonzalez, 2017; Xie and Zhang, 2014). Among them, miR-185 has been revealed as a tumor inhibitor in several cancers such as nasopharyngeal carcinoma (Liu and Li, 2017), non-small cell lung cancer (NSCLC) (Li and Ma, 2015) and OC (Xiang and Ma, 2014). Importantly, miR-185 was demonstrated to negatively target the 3’□untranslated region (3’-UTR) of SRY-box 9 (SOX9) in a former study (Lei and Shi, 2018). Aberrant expression of SOX9 has been suggested participating in the pathogenesis of several cancers (Lu and Fang, 2008; Thomsen and Ambroisine, 2010). Likewise, abnormal SOX9 expression has been demonstrated to promote OC cell survival in hypoxic condition and to affect OC aggressiveness (Raspaglio and Petrillo, 2014). Meanwhile, a previous study has identified that SOX9 could trigger Wnt/β-catenin pathway activation in prostate cancer (Ma and Ye, 2016). It is well-known that the Wnt/β-catenin signaling pathway plays critical roles in cancer development (Li and Zhang, 2015). A recent study suggested that inhibition of Wnt/β-catenin could suppress stemness and drug resistance of OC cells (Ruan and Liu, 2019). However, the effects of DEX and the cytokines above on OC remain unknown. Hence, this study was designed to explore the roles of DEX and these potential biomarkers in OC development.

## Materials and methods

### Ethics statement

The study was approved by the Clinical Ethical Committee of Guangzhou Medical University. All experimental procedures were performed in line with the ethical guidelines for the study of experimental pain in conscious animals.

### Cell culture

Human OC cell lines SKOV3 and HO-8910 (purchased from Institute of Cell Biology, Chinese Academy of Sciences, Shanghai, China) were cultured in a 37□ constant incubator with 5% CO_2_, and the culture solution was refreshed everyday. Cells were then passaged when the cell confluence reached 80%.

### Cell transfection and grouping

The well-grown SKOV3 and HO-8910 cells at passage 3 were randomly assigned into 4 groups, which were respectively treated with 0 nmol/L, 1 nmol/L, 10 nmol/L and 100 nmol/L DEX to measure the effect of different doses of DEX on cell growth and to select the best dose of DEX for the following transfection.

Cell transfection: the SKOV3 and HO-8910 cells were treated with an appropriate dose of DEX for 24 h. After that, the SKOV3 cells were randomized into DEX + negative control (NC) group (transfected with mimics NC) and DEX + miR-185 group (transfected with miR-185 mimics), while the HO-8910 cells were assigned into DEX + NC group (transfected with antisense NC), DEX + anti-miR-185 group (transfected with miR-185 antisense) and DEX + anti-miR-185 + si-SOX9 group (transfected with miR-185 antisense and siRNA against SOX9). The transfections were performed as per the instructions of a Lipofectamine™ 2000 kit (Invitrogen Inc., Carlsbad, CA, USA). The transfection efficiencies were assessed with reverse transcription-quantitative polymerase chain reaction (RT-qPCR) 24 h following transfection.

### 3-(4, 5-dimethylthiazol-2-yl)-2, 5-diphenyltetrazolium bromide (MTT) assay

Cells in each group were seeded onto 96-well plates (1 × 10^4^ cells/well) and then incubated at 37□ with 5% CO_2_ for 24 h, and 6 duplicated wells were set for each group. Thereafter, each well was treated with 5 mg/mL MTT solution (Sigma-Aldrich Chemical Company, St Louis, MO, USA) and the cells were cultured for an additional 4 h. A total of 200 μL dimethyl sulfoxide was added after the supernatant was discarded. Following 10 min of vibration, the optical density (OD) value at 490 nm of each well was evaluated using a microplate reader, and the average value of 6 wells was recorded.

### 5-ethynyl-2’-deoxyuridine (EdU) labeling assay

In accordance with the instructions of an EdU assay kit purchased from Guangzhou RiboBio Co., Ltd (Guangzhou, Guangdong, China), cells were seeded onto 48-well plates (8000 cells/well) and incubated with 20 μmol/L EdU for 24. Followed by 30 min of 4% polyformaldehyde fixing and 10 min of 0.5% Triton X-100 permeabilization, each well of cells were stained with 150 μL reaction solution for 30 min and 150 μL 1 × Hochest33258 for 5 min. A fluorescence microscope (Olympus Optical Co., Ltd, Tokyo, Japan) was utilized to observe the cells. Five areas were randomly selected in which the blue fluorescence refers all cells while the red fluorescence indicates the replicating cells influenced by EdU, and then the EdU-positive cell rates were calculated.

### Colony formation assay

The well-grown cells at passage 3 were detached using 0.025% trypsin and incubated in 6-well plates (1000 cell/well) at 37□ with 5% CO_2_ for 2 w, after which they were treated with 75% methanol for 30 min, stained with 0.5% crystal violet for 15 min and finally observed under an optical microscope (IX51, Olympus Optical Co., Ltd, Tokyo, Japan) to analyze and calculate the number of cell colonies. Each experiment was performed for 3 times.

### Flow cytometry

An apoptosis assay was conducted using an Annexin V-fluorescein isothiocyanate (FITC)/propidium iodide (PI) cell apoptosis kit (KeyGen, Nanjing, Jiangsu, China). Briefly, cells in each group were sorted onto 6-well plates at the density of 2 × 10^6^ cells per well, after which they were stained with the mixture of 5 μL ALexa Flour 488 AnnexinV-FITC and 1 μL PI (100 mg/mL) and then incubated at room temperature in the dark for 15 min. Next, each well of cells were treated with 400 μL 1 × Annexin binding buffer and then the cell apoptosis was detected using a flow cytometer. Approximately 1 × 10^4^ cells were included and the percentage of Annexin V+ (apoptotic) cells in each group was evaluated.

### Terminal deoxynucleotidyl transferase (TdT)-mediated dUTP nick end labeling (TUNEL) staining

Cell suspension was sorted into confocal dishes (Thermo Fisher Scientific Inc., Waltham, MA, USA), which was further cultured in a 37□ incubator with 50 mL/L CO_2_. When the cell confluence got to 80%, the cells were subjected to TUNEL staining as per the kit instructions (R&D Systems Inc., Minneapolis, MN, USA). Followed by 4’, 6-diamidino-2-phenylindole (DAPI) (10 μg/mL) staining for 10 min, the cell apoptosis was observed under a confocal microscope (Radiance 2100, Bio-Rad Laboratories Inc., Hercules, CA, USA).

### Transwell assay

Cells were seeded into the Transwell chambers at the density of 1 × 10^5^ cells/mL, and the apical chambers which coated with Matrigel were treated with serum-free media containing different doses of DEX (0 nmol/L, 1 nmol/L, 10 nmol/L and 100 nmol/L) while the basolateral chambers were treated with normal media. After 24 h of culture, cells invading to the basolateral chamber were stained using crystal violet and observed under a microscope.

### Scratch test

Briefly, 5 parallel guide lines were produced at the back side of 12-well plates. SKOV3 and HO-8910 cells were plated into 12-well plates at the density of 1 × 10^5^cells/mL. A 10 μL pipette tip was applied to produce scratches on the adherent cells perpendicular to the back guide lines, and then the suspension cells were rinsed with binding buffer. Next, the cells were cultured in serum-free media containing different doses of DEX (0 nmol/L, 1 nmol/L, 10 nmol/L and 100 nmol/L), and the cell growths at 0 h and 24 h were recorded for wound-healing rate calculation with the formula as follows: wound-healing rate (%) = (scratch width at 0 h - width at 24 h)/width at 0 h.

### RT-qPCR

Total RNA of cells were extracted using Trizol one-step method (Invitrogen, Carlsbad, CA, USA) and the ultraviolet analysis and formaldehyde denaturing gel electrophoresis were applied to identify the high-quality RNA. After that, l μg RNA was collected and reverse-transcribed into cDNA by avian myeloblastosis virus reverse transcriptase. qPCR was conducted using a SYBR Green assay, and the PCR primers were designed and synthesized by Shanghai Sangon Biotech Co., Ltd (Shanghai, China) (Table I), with U6 or β-actin was set as the internal reference. The PCR reaction system was consisted of 1.0 μL cDNA, 10 μL 2× SYBR Green Mix, 0.5 μL Forward primer (10 μM), 0.5 μL reverse primer (10 μM) and RNase free water until the solution arrived at 20 μL. The PCR reaction condition was as follows: pre-denaturation at 94°C for 5 min, followed by 40 cycles of denaturation at 94°C for 40 s, annealing at 60°C for 40 s, extension at 72°C for 1 min, and a final extension at 72°C for 10 min. The production was identified using agarose gel electrophoresis. Data was analyzed using 2^-ΔΔCt^ method in which 2^-ΔΔCt^ refers to the ratio of the target gene expression between the experimental and control groups, the formula was as follows: ΔΔCt = [Ct (target gene) - Ct (internal control gene)] _experimental group_ - [Ct (target gene) - Ct (internal control gene)] _control group_.

**Table I.**
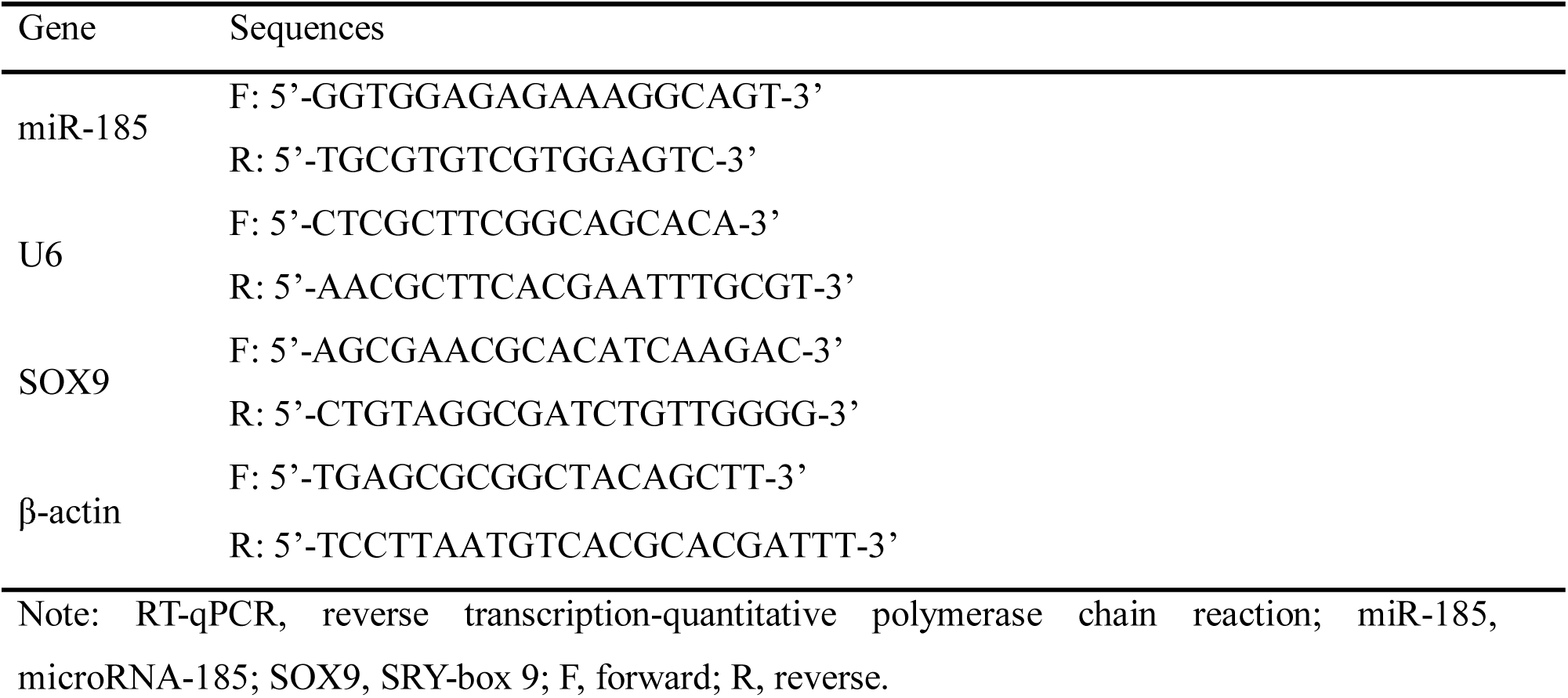
Primer sequences for RT-qPCR

### Western blot analysis

Proteins were extracted from cells and their concentrations were evaluated as per the instructions of a bicinchoninic acid assay kit (Wuhan Boster Biological Technology, Ltd., Hubei, China). Followed by blending with the loading buffer, the extracted proteins were boiled at 95°C for 10 min and then loaded into the wells with 30 µg per well. Next, the proteins were detached using 10% sodium dodecyl sulfate-polyacrylamide gel electrophoresis (Wuhan Boster Biological Technology, Ltd., Hubei, China) with the voltage from 80 V to 120 V and transferred to polyvinylidene difluoride membranes (100 mV for 45 ∼ 70 min). Thereafter, the membranes were sealed with 5% bovine serum albumin at room temperature for 1 h and incubated with the following antibodies at 4□ overnight: cleaved caspase-3 (1 µg/mL, ab2302), Bax (1:1000, ab53154), Bcl-2 (1:1000, ab32124), SOX9 (1:1000, ab185230), β-catenin (1:2000, ab32572), C-myc (1:1000, ab32072) and cyclinD1 (1:200, ab16663). All these antibodies were purchased from Abcam Inc., (Cambridge, MA, USA). Followed by Tris-buffered saline with Tween 20 (TBST) washing (3 times/5 min), the protein were incubated with secondary antibodies (ZSGB-Bio, Beijing, China) for 1 h and washed with TBST for 3 times before chemiluminescence developing. A Gel Doc EZ Imager (Bio-Rad, California, USA) was used to develop the protein bands. The grey value analysis for target bands was performed using Image J software (National Institutes of Health. Bethesda, Maryland, USA).

### Dual luciferase reporter gene assay

An online miRNA target detection program was conducted to predict the binding sites of miR-185 and SOX9 (miRNA.org). The 3’ untranslated region (3’UTR) sequence of SOX9 which containing the miR-185 binding site was synthesized, and then a SOX9 3’UTR wild-type (WT) plasmid (SOX9-WT) and another SOX9 3’UTR mutant-type plasmid (SOX9-MUT) plasmid were constructed. Well-designed plasmids along with miR-185 mimic or mimic NC were co-transfected into 293T cells. Cells were collected and lysed 48 h after transfection, and their luciferase activities were detected based on luciferase assay kit (BioVision, SanFrancisco, CA, USA) and Glomax20/20 illuminometer (Promega, Madison, Wisconsin, USA).

### Immunofluorescence assay

Cells were sorted into 6-well plates which laid with 0.1 gelatin-coated cover glass until 80% ∼ 90% of cell confluence was reached, the culture solution was discarded after which the cells were fixed with 4% polyformaldehyde for 20 min. Followed by 10% goat serum sealing for 1 h, the cells were incubated with primary rabbit polyclonal antibodies SOX9 and β-catenin, fluorescence-labeled secondary antibody and stained with 4’, 6-diamidino-2-phenylindole at room temperature for 30 min, after which the staining was observed under a confocal microscope.

### Xenograft tumors in nude mice

BALB/c nude mice (male, 16-18 g, 4-6 weeks) purchased from Beijing Vital River Laboratory Animal Technology Co., Ltd., (Beijing, China) were fed in a specific pathogen free grade animal center. After hypodermic injection of 100 μL (1 × 10^6^) cells into the nude mice, the length (L) and width (W) of the tumors were recorded every 5 days, and the tumor volume (V) was calculated as follows: V (mm^3^) = L × W^2^ × 0.5, after which a tumor growth curve was produced. The mice were euthanized at day 30, and the subcutaneous tumors were extracted and weighted.

### Immunohistochemistry

After weighted, the tumors were successively fixed with 4% polyformaldehyde for 24 h, dehydrated in series of alcohol, waxed twice, embedded in paraffin and cut into 4 μm sections. The paraffin-embedded sections were set on cover glasses coated with polylysine, baked for 2 h and then cooled at room temperature. Next, the sections were dewaxed in xylene and ethanol and added into 0.01 mol/L citrate buffer for 15 min of high pressure thermal treatment. After cooled at room temperature, the sections were treated with 3% H_2_O_2_ for 15 min to block the activity of endogenous peroxidase, which was followed by phosphate-buffered saline (PBS) washing for 15 min and incubated with primary antibodies at 4□ overnight. The second day, the sections were washed in PBS for 15 min and incubated with corresponding secondary antibody at 37□ for 40 min, which was followed by PBS washing for 15 min again. Then the sections were successively stained with 2, 4-diaminobutyric acid, rinsed with water for 15 min, counterstained with hematoxylin for 1 min, differentiated, turned to blue, washed, dehydrated and sealed. A NC was set with the primary antibodies were replaced as PBS, and the positive sections provided by the company were set as positive control, and the sections were observed under a microscope. Cells with positive ki67, cyclinD1 and p53 were showing with yellow or brown cytoplasm. Followed by comprehensive observations under low magnification, the integrated OD (IOD) values of cells were detected using an Image-propluse6.0 program (Media Cybernetics, Silver Spring, MD, USA) under a × 200 microscope magnification. A total of 5 views were randomly selected and the average IOD value was calculated.

### Hematoxylin-eosin (HE) staining

The xenograft tumors were embedded with paraffin and sectioned, which was followed by HE staining and pathomorphologic characteristics observation under a microscope.

### Statistical analysis

The Statistical Package for the Social Sciences (SPSS) 21.0 (IBM Corp. Armonk, NY, USA) was applied for data analysis. The normality test was conducted using the Kolmogorov-Smirnov method. Measurement data were showing as normal distribution and expressed as mean ± standard deviation. Differences between pairwise groups were evaluated using the *t*-test while the differences in multiple groups were compared using one-way analysis of variance (ANOVA). *p* was two-sided and *p* < 0.05 was considered to show a statistical significance.

## Results

### DEX inhibits growth, metastasis and resistance to apoptosis of OC cells

The viabilities of OC cells were evaluated using MTT assay, and the results suggested that the viabilities of SKOV3 and HO-8910 both decreased after increased doses of DEX treatment, which was showing with time- and dose-dependence (Figure 1A), indicating that DEX could suppress the proliferation of OC cells. Meanwhile, the number of cell colonies and EdU-positive cell rate both showed DEX dose-dependent declines after increased doses of DEX treatment (Figure 1B-C). Moreover, the apoptotic rate of SKOV3 and HO-8910 cells significantly elevated as DEX dose increased (Figure 1D-E). The expression of apoptosis-related proteins cleaved caspase3, and Bax enhanced while the expression of Bcl-2 notably reduced after increased doses of DEX treatment (*p* < 0.05) (Figure 1F). Besides, according to Transwell assay and scratch test, the invasion and migration abilities of SKOV3 and HO-8910 cells significantly reduced after an ascending series of DEX treatment (Figure 1G-H). The results above indicated that DEX has a dose-dependent inhibitory effect on OC cell growth, metastasis and resistance to apoptosis.

**Figure 1.**
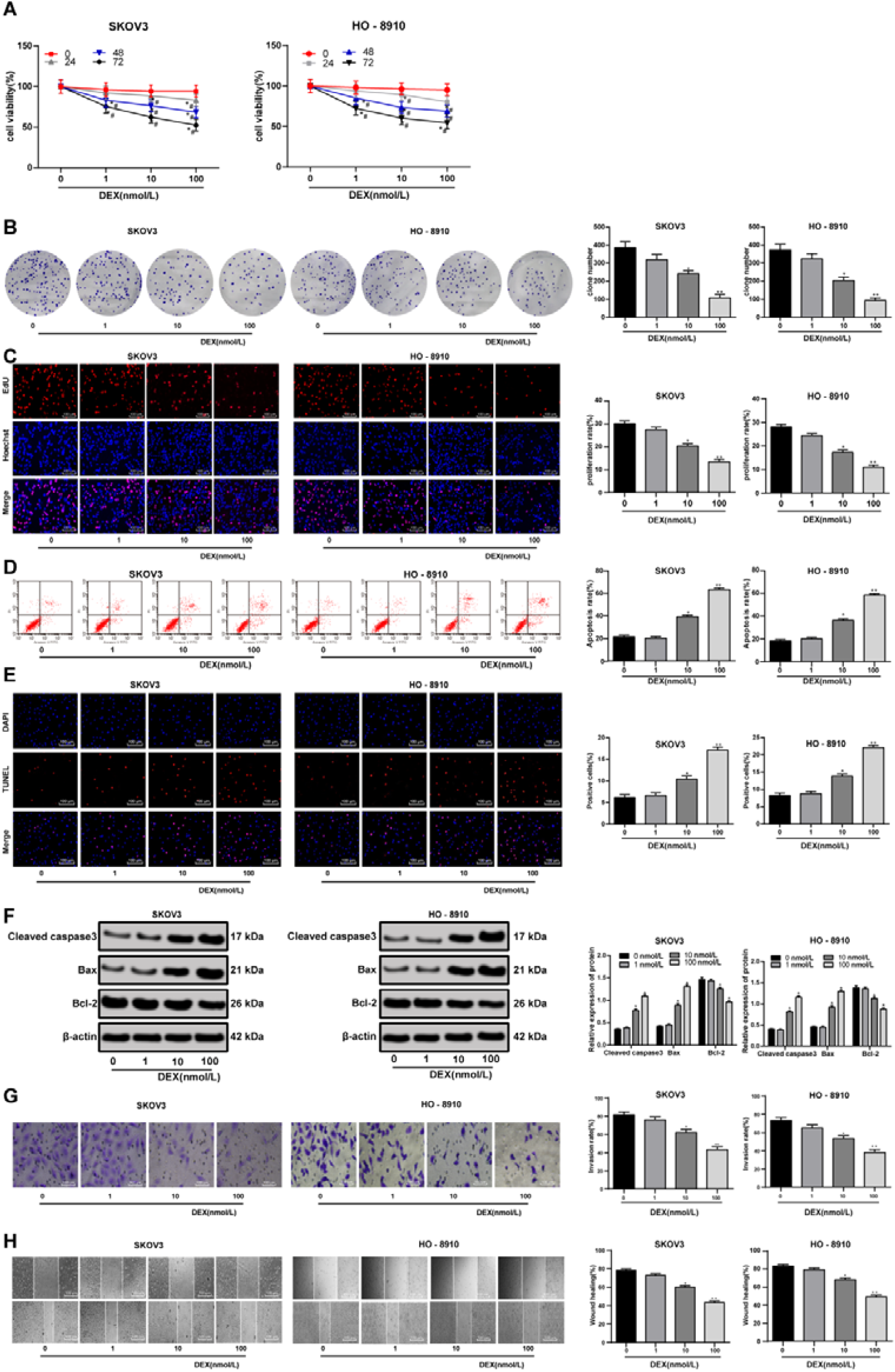
DEX has a dose-dependent inhibitory effect on OC growth. A, cell viability detected using MTT assay; B, number of cell colonies detected using colony formation assay; C, DNA replication measured using EdU assay; D, cell apoptosis detected using flow cytometry; E, cell apoptosis detected using TUNEL assay; F, expression of apoptosis-related proteins detected using Western blot analysis; G, cell invasion evaluated using Transwell assay; H, cell migration detected using scratch test; all experiments were performed for three times, and one-way ANOVA was used to determine statistical significance; * compared to 0 nmol/L DEX treatment, *, *p* < 0.05; **, *p* < 0.01; DEX, dexmedetomidine; OC, ovarian cancer; MTT, 3-(4, 5-dimethylthiazol-2-yl)-2, 5-diphenyltetrazolium bromide; EdU, 5-ethynyl-2’-deoxyuridine labeling; TUNEL, terminal deoxynucleotidyl transferase (TdT)-mediated dUTP nick end labeling; ANOVA, analysis of variance.

### DEX up-regulates miR-185 expression but suppresses SOX9, β-catenin, C-myc and cyclinD1 expression in OC cells

The antioxidative, anti-inflammatory properties of DEX have been well established (Wang and Zhao, 2015). Over expression of miR-185 has been found to suppress proliferation and to promote apoptosis of OC cells (Xiang, 2014). Meanwhile, a former study has revealed that over-expressed SOX9 and β-catenin were implicated in the development of OC development (Xiao and Li, 2019). In the current study, it was shown that DEX has a dose-dependent promoting effect on miR-185 expression but inhibitory effects on SOX9, β-catenin, C-myc and cyclinD1 expression in OS cells (all *p* < 0.05) (Figure 2A-B).

**Figure 2.**
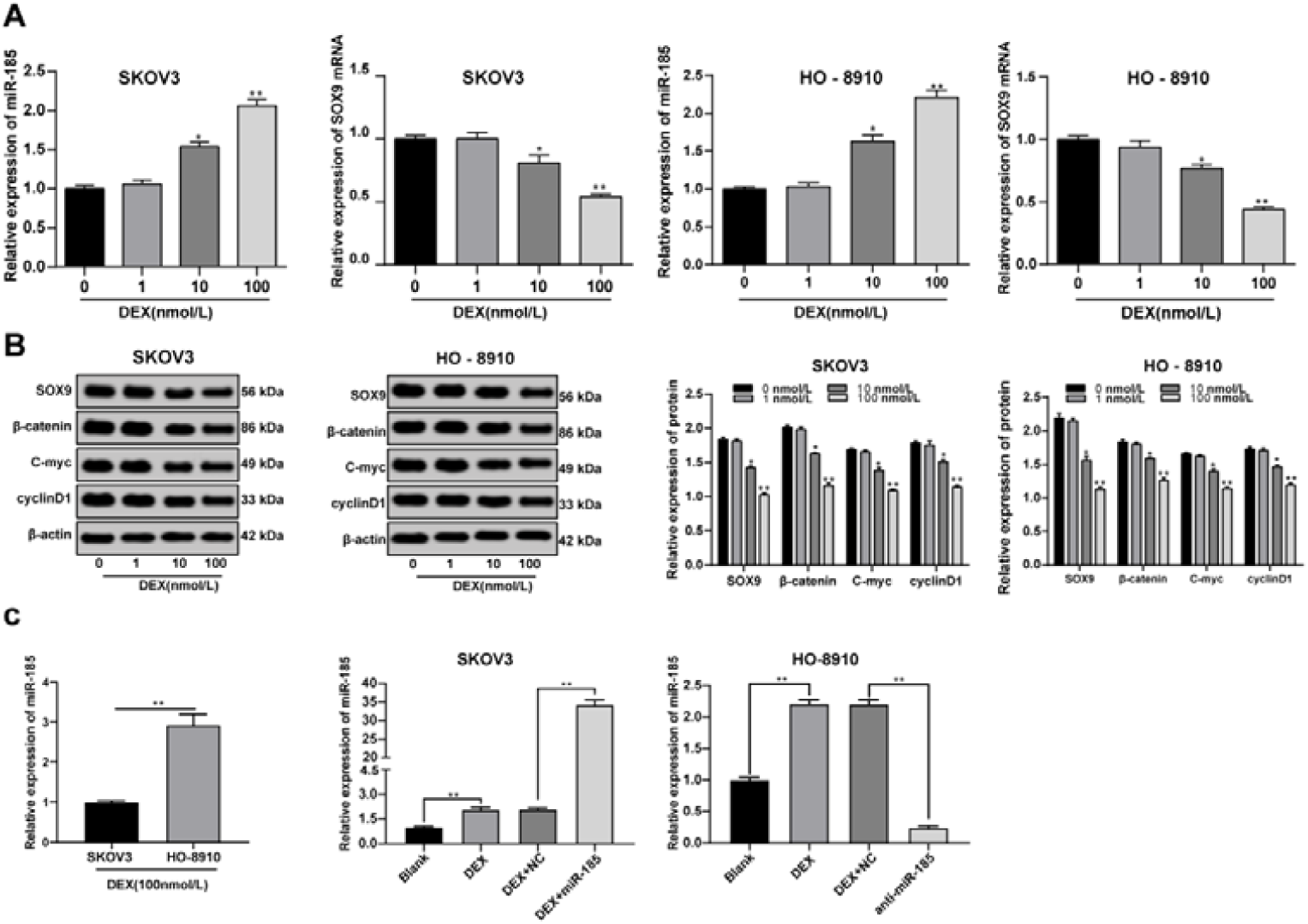
DEX up-regulates miR-185 expression but suppresses SOX, β-catenin, C-myc and cyclinD1 expression in OC cells. A, miR-185 and SOX9 mRNA expression detected using RT-qPCR; protein expression of SOX9, β-catenin, C-myc and cyclinD1 measured using Western blot analysis; C, miR-185 expression in SKOV3 and HO-8910 cells after transfection; all experiments were performed for three times, and one-way ANOVA was used to determine statistical significance; * compared to 0 nmol/L DEX treatment or compared to cells without miR-185 transfection, *, *p* < 0.05; **, *p* < 0.01; DEX, dexmedetomidine; miR-185, microRNA-185; SOX9, SRY-box 9; OC, ovarian cancer; RT-qPCR, reverse transcription-quantitative polymerase chain reaction; ANOVA, analysis of variance.

Following the results that DEX inhibited OC cell proliferation and the evidence that DEX enhanced miR-185 expression in OC cells, the mechanisms of miR-185 expression in DEX-treated OC cells needed further investigation. Thus, the SKOV3 cells were transfected with miR-185 mimic and the HO-8910 cells were transfected with miR-185 antisense after 100 nmol/L DEX treatment for comparative studies. The results of RT-qPCR suggested miR-185 was over-expressed in SKOV3 cells while lowly-expressed in HO-8910 cells (*p* < 0.05) (Figure 2C), which indicated that the plasmids were successfully transfected.

### DEX and over-expressed miR-185 inhibits OC cell growth and metastasis

After 100 nmol/L DEX treatment and corresponding transfection, the viability of the SKOV3 cells (with over-expressed miR-185) notably reduced, while that of the HO-8910 cells (with lowly-expressed miR-185) significantly enhanced (*p* < 0.05) (Figure 3A-C). Similarly, the apoptotic rate of the SKOV3 cells obviously elevated but that of the HO-8910 cells notably decreased (*p* < 0.05) (Figure 3D-E). Meanwhile, the cleaved caspase3 and Bax levels significantly increased while the Bcl-2 level decreased in the SKOV3 cells with over-expression of miR-185, while those in the HO-8910 cells presented an opposite trend (*p* < 0.05) (Figure 3F). Likewise, the invasion and migration of the SKOV3 cells were significantly suppressed while those of the HO-8910 cells were notably enhanced (*p* < 0.05) (Figure 3G-H). These results indicated that down-regulation of miR-185 could reverse the inhibitory effect of DEX on OC cells.

**Figure 3.**
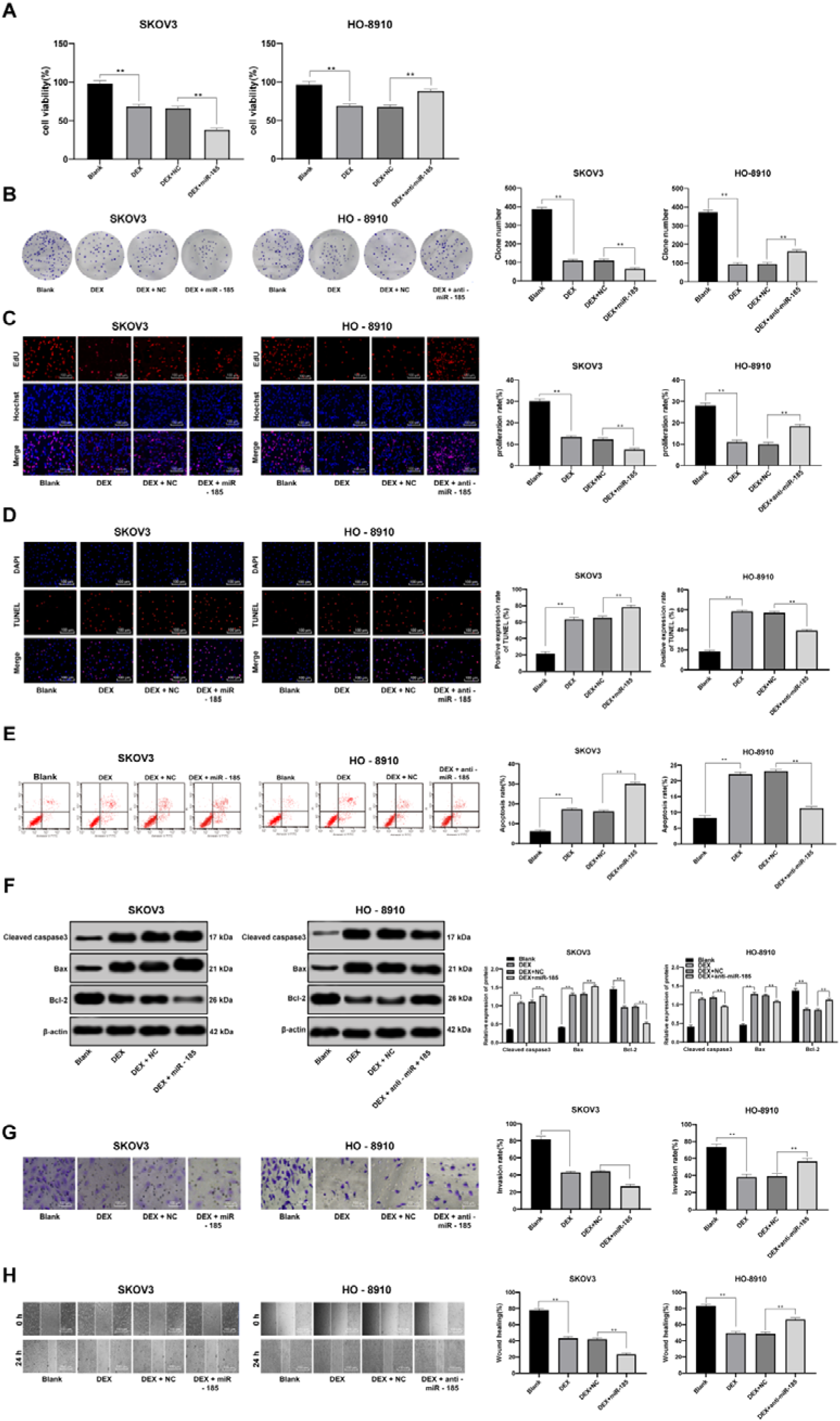
DEX and over-expressed miR-185 inhibits OC cell growth and metastasis. A, cell viability detected using MTT assay; B, number of cell colonies detected using colony formation assay; C, DNA replication measured using EdU assay; D, cell apoptosis detected using flow cytometry; E, cell apoptosis detected using TUNEL assay; F, expression of apoptosis-related proteins detected using Western blot analysis; G, cell invasion evaluated using Transwell assay; H, cell migration detected using scratch test; all experiments were performed for three times, and one-way ANOVA was used to determine statistical significance; * compared to cells without miR-185 transfection, *, *p* < 0.05; **, *p* < 0.01; DEX, dexmedetomidine; miR-185, microRNA-185; OC, ovarian cancer; MTT, 3-(4, 5-dimethylthiazol-2-yl)-2, 5-diphenyltetrazolium bromide; EdU, 5-ethynyl-2’-deoxyuridine labeling; TUNEL, terminal deoxynucleotidyl transferase (TdT)-mediated dUTP nick end labeling; ANOVA, analysis of variance.

### miR-185 inhibits SOX9 expression in DEX-treated OC cells

Based on the evidence confirmed above that the miR-185 expression was promoted while the SOX9 expression was inhibited after DEX treatment, the correlations between miR-185 and SOX9 were further explored. The computer-based program (miRNA.org) suggested that miR-185 could directly bind to the 3□UTR of SOX9 (Figure 4A) Moreover, the dual luciferase reporter gene assay results suggested that the luciferase activity of cells co-transfected with SOX9 (WT) and miR-185 significantly reduced, which further identified the target relations between miR-185 and SOX9 (*p* < 0.05) (Figure 4B). In addition, the SOX9 expression in SKOV3 and HO-8910 cells after 100 nmol/L DEX treatment and corresponding transfection was measured. The results showed that the SOX9 expression in the SKOV3 cells notably declined while that in the HO-8910 cells obviously enhanced (*p* < 0.05) (Figure 4C-D). These results suggested that miR-185 negatively targets SOX9 expression.

**Figure 4.**
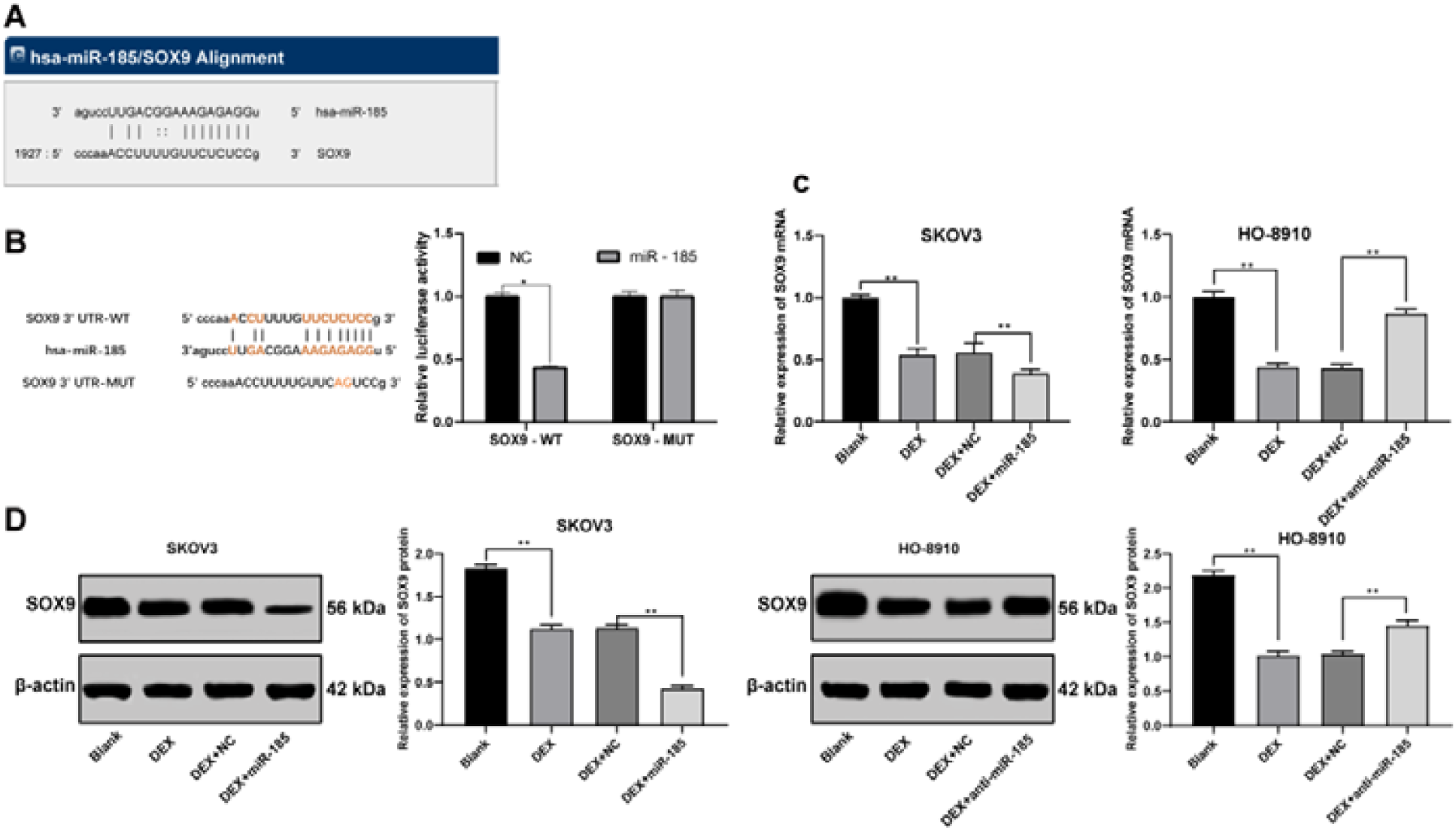
miR-185 directly binds to SOX9. A, target relation between miR-185 and SOX9 predicted via miRNA.org program; B, target relation between miR-185 and SOX9 identified using dual luciferase reporter gene assay; C, SOX9 mRNA expression in SKOV3 and HO-8910 cells detected using RT-qPCR; D, SOX9 protein expression in SKOV3 and HO-8910 cells after detected using Western blot analysis; in panel B, the experiment was performed for 3 times and *t* test was used to analyze statistical significance; in panel C and D, all experiments were performed for 3 times and one-way ANOVA was used to determine statistical significance; * compared to 100 nmol/L DEX treatment only, *, *p* < 0.05; miR-185, microRNA-185; SOX9, SRY-box 9; RT-qPCR, reverse transcription-quantitative polymerase chain reaction; DEX, dexmedetomidine; ANOVA, analysis of variance.

### SOX9 knockdown inhibits OC cells proliferation and metastasis

Lowly-expressed miR-185-promoted HO-8910 cell growth trigged us to study whether SOX9 expression was implicated in these events, thus a rescue experiment was performed to explore the roles of SOX9 in OC progression. It was shown that even when miR-185 expression was inhibited, SOX9 knockdown led to significant decline in HO-8910 cell viability, number of cell colonies and EdU-positive cell rates (all *p* < 0.05) (Figure 5A-C), and the cell apoptotic rate and cleaved caspase 3 and Bax expression significantly elevated while the Bcl-2 expression notably decreased (all *p* < 0.05) (Figure 5D-F). Besides, the invasion and migration of HO-8910 cells obviously reduced after SOX9 knockdown (all *p* < 0.05) (Figure 5G-H). These results demonstrated that SOX9 knockdown could reverse the promoting effect of low-expression of miR-185-induced OC cell proliferation and resistance to apoptosis.

**Figure 5.**
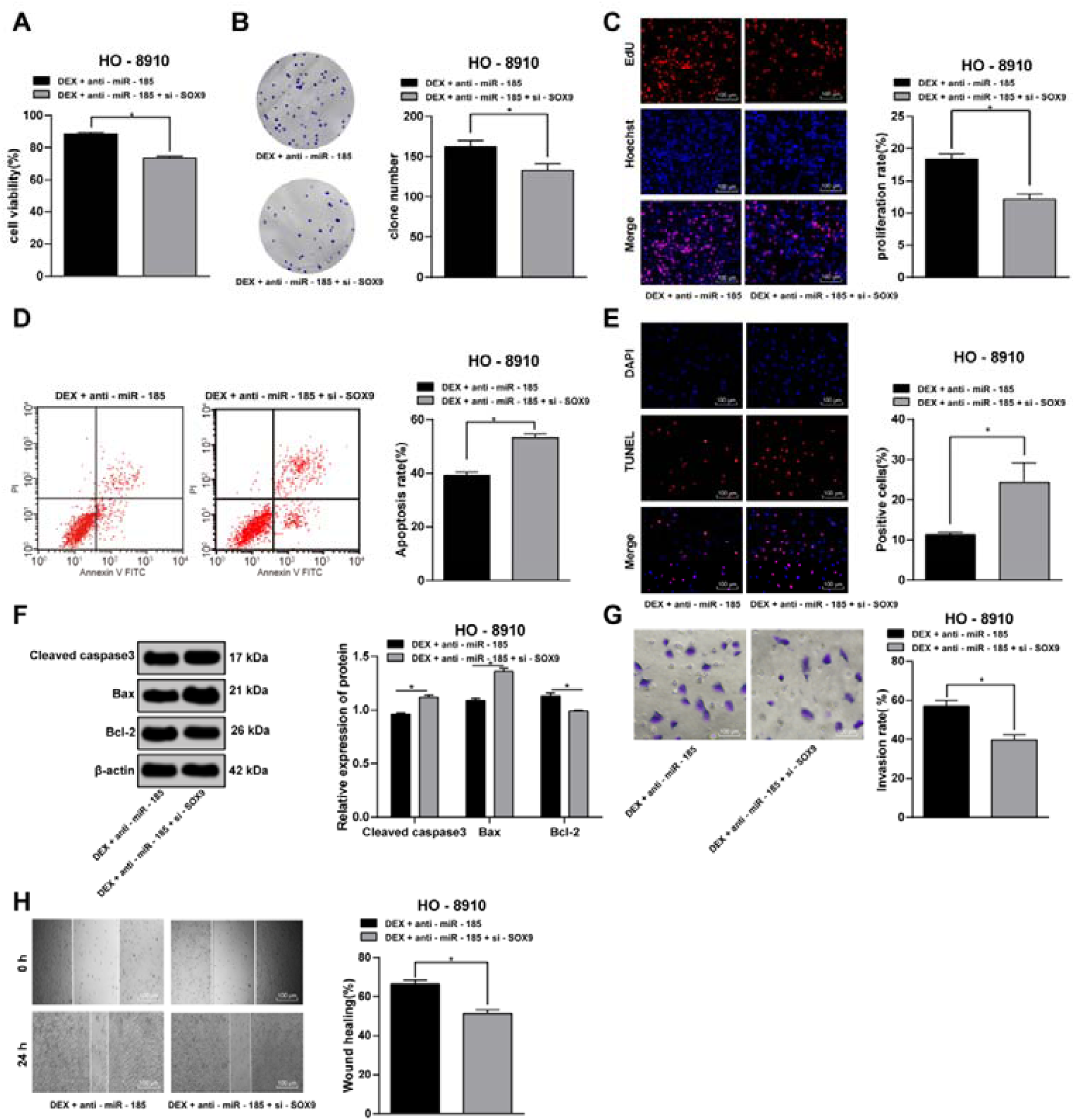
SOX9 knockdown inhibits OC cells proliferation and metastasis. A, cell viability detected using MTT assay; B, number of cell colonies detected using colony formation assay; C, DNA replication measured using EdU assay; D, cell apoptosis detected using flow cytometry; E, cell apoptosis detected using TUNEL assay; F, expression of apoptosis-related proteins detected using Western blot analysis; G, cell invasion evaluated using Transwell assay; H, cell migration detected using scratch test; all experiments were performed for 3 times and *t* test was used to analyze statistical significance * compared to cells without SOX9 knockdown treatment, *, *p* < 0.05; **, *p* < 0.01; SOX9, SRY-box 9; OC, ovarian cancer; MTT, 3-(4, 5-dimethylthiazol-2-yl)-2, 5-diphenyltetrazolium bromide; EdU, 5-ethynyl-2’-deoxyuridine labeling; TUNEL, terminal deoxynucleotidyl transferase (TdT)-mediated dUTP nick end labeling.

### DEX up-regulates miR-185 expression to inactivate the SOX9/Wnt/β-catenin signaling pathway

Based on the results above that DEX could promote miR-185 expression thus negatively targeting SOX9 expression, we further investigate the role of miR-185 and SOX9 in the protein levels of β-catenin, C-myc and cyclinD1 in OC cells. The results suggested that after DEX treatment and corresponding transfection, the protein levels of SOX9, β-catenin, C-myc and cyclinD1 significantly reduced in the SKOV3 cells, while those in the HO-8910 cells presented an opposite trend (all *p* < 0.05) (Figure 6A). Moreover, the fluorescence intensities of SOX9 and β-catenin in the SKOV3 cells were notably lower than those in the HO-8910 cells, likewise, the HO-8910 cells with SOX9 knockdown presented reduced fluorescence intensities of SOX9 and β-catenin (all *p* < 0.05) (Figure 6B).

**Figure 6.**
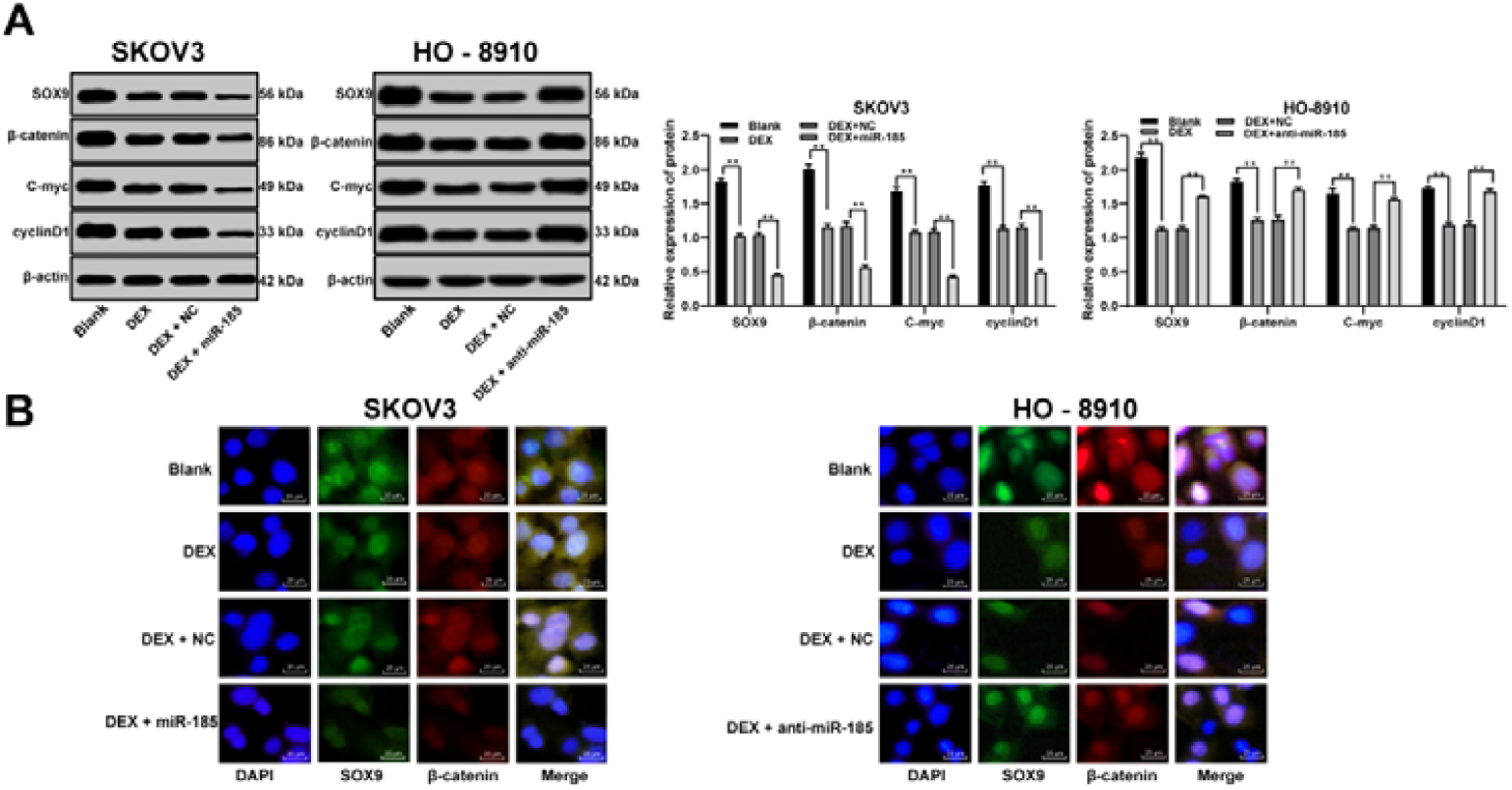
DEX up-regulates miR-185 expression to inactivate the SOX9/Wnt/β-catenin signaling pathway. A, protein expression of SOX9, β-catenin, C-myc and cyclinD1 detected using Western blot analysis; B, fluorescence intensities of SOX9 and β-catenin evaluated using immunofluorescence assay; in panel A, the experiment was performed for 3 times and one-way ANOVA was used to determine statistical significance; * compared to 100 nmol/L DEX treatment only, *, *p* < 0.05; DEX, dexmedetomidine; miR-185, microRNA-185; SOX9, SRY-box 9; ANOVA, analysis of variance.

### DEX and over-expressed miR-185 suppresses OC tumor growth in vivo

Following the *in vitro* studies concluded above, a xenograft tumor model was established to explore the role of DEX and miR-185 *in vivo*. After DEX treatment and corresponding miR-185 transfection, the SKOV3 and HO-8910 cells were injected into the nude mice and the volume and weight of tumors were recorded. It was shown that the weight and volume of tumors in mice injected with SKOV3 cells were notably decreased while those in mice injected with HO-8910 cells were obviously increased (all *p* < 0.05) (Figure 7A-B). Moreover, the immunohistochemistry results of ki67, cyclinD1 and p53 suggested that the SKOV3 transfection led to significantly declined positive ki67, cyclinD1 and p53 in tumor cells, while the HO-8910 transfection contributed to an opposite trend (all *p* < 0.05) (Figure 7C). Besides, HE staining results suggested that the tumor tissues in mice transfected with SKOV3 cells presented obvious necrosis, while those in mice transfected with HO-8910 cells presented typical tumor morphologies with diverse cell sizes and shapes, large nucleus, frequent mitosis, little karyopyknosis and necrosis (Figure 7D). These results suggested DEX and over-expression of miR-185 could suppress OC tumor growth *in vivo*.

**Figure 7.**
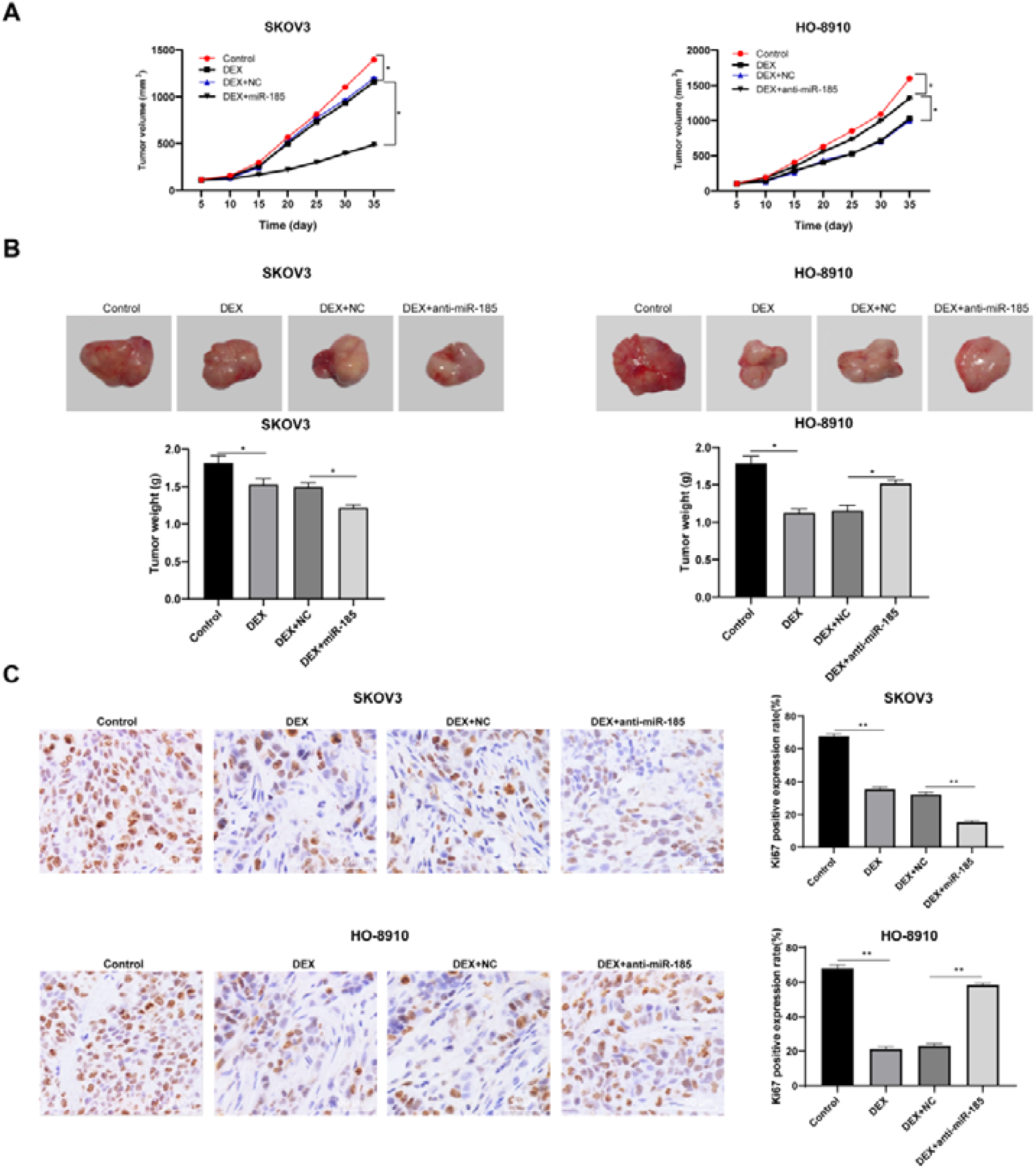
DEX and over-expressed miR-185 suppresses OC tumor growth *in vivo*. A, tumor volume detection after OC cell transfection; B, tumor weight detection after OC cell transfection; C, ki67, cyclinD1 and p53 expression detected using immunohistochemistry; D, morphologies of tumor tissues observed using HE staining; in panel A, B and C, all experiments were performed for 3 times and one-way ANOVA was used to determine statistical significance;* compared to 100 nmol/L DEX treatment only, *, *p* < 0.05; miR-185, microRNA-185; OC, ovarian cancer; HE, hematoxylin and eosin; DEX, dexmedetomidine; ANOVA, analysis of variance.

## Discussion

OC is a life-threatening malignancy which has not seen a major therapeutic improvement over 3 decades (Basuli and Tesfay, 2017). DEX is a widely used anesthetic and also has organ-protective (Heimisdottir, 2016), neuroprotective (Shanbehzadeh and Salavati, 2018) and in some cases, has anticancer (Wang and Xu, 2018) effects. Meanwhile, dysregulation of miRNAs has been well-established involving in a number of diseases including cancers (Palanichamy and Rao, 2014). The current study was designed to figure out the possible relationship between DEX and miR-185 and their effects on OC development, with a conclusion was drawn that DEX could inhibit OC development via up-regulating miR-185 expression and inactivating the SOX9/Wnt/β-catenin signaling pathway.

Initially, our study found that DEX showed a dose-dependent inhibitory effect on OC growth, invasion and migration. Emerging studies have suggested that anesthetics may also have anti-tumor properties such as morphine (Yam and He, 2011) and propofol (Xu and Du, 2013). Likewise, DEX has been demonstrated to suppress cell proliferation and migration and to promote cell death in osteosarcoma cells (Wang, 2018). Besides, a previous study also suggested that DEX treatment could inhibit OC growth and the tumor weight of OS rats *in vivo* (Cai and Tang, 2017). Recently, Lei Hou *et al*. found that DEX could improve the immune function of OC rats and inhibit the invasion and migration of OC cells via inactivating the insulin-like growth factor 2 signaling pathway (Tian and Hou, 2019). Meanwhile, in this present study, DEX also presented a dose-dependent promoting role in OC cell apoptosis, with increased expression of cleaved caspase 3 and Bax while declined expression of Bcl-2 shown after DEX treatment. Caspase 3 and Bax/Bcl-2 ratio are well-known proapoptotic factors (Freitas and Alves, 2011). Bcl-2 is an antiapoptotic protein which regulates apoptosis via modulating the permeability of mitochondrial membrane, while Bax is capable of damaging the outer mitochondrial membrane (Lv and Liang, 2018; Peng and Chen, 2016). Thus, DEX could inhibit proliferation, metastasis and resistance to apoptosis of OC cells.

To further identify the molecular mechanism participating in the inhibitory effect of DEX on OC development, we detected the miR-185 expression in OC cells after DEX treatment. Importantly, DEX elevated miR-185 expression in OC cells but down-regulated SOX9 expression. Meanwhile, according to bioinformation prediction and dual luciferase reporter gene assay, miR-185 was identified to negatively bind to 3’-UTR of SOX9, which is coincidence with the study published by Zhengwen Lei *et al*. (Lei, 2018). In our study, both over-expression of miR-185 in SKOV3 cells or knockdown of SOX9 in OC cells inhibited cell growth and metastasis. Emerging studies have revealed that miR-185 works as a tumor suppressor. For instance, miR-185 could inhibit the proliferative potential of human colorectal cancer cells (Liu and Lang, 2011). JS Imam *et al*. suggested that miR-185 is lowly expressed in several human cancers including OC and it could impede tumor growth and progression (Imam and Buddavarapu, 2010). SOX9 is a key transcription factor required for cancer development whose overexpression promotes cell proliferation and growth in lung cancer (Jiang and Fang, 2010). Likewise, overexpression of SOX9 has been demonstrated to cause neoplasia in murine prostate and to promote tumor invasion (Cai and Wang, 2013). Furthermore, our study found that DEX treatment or overexpression of miR-185 reduced the protein expression of SOX9, β-catenin, C-myc and cyclin D1 in OC cells. C-myc and cyclinD1 are two major downstream target of Wnt/β-catenin signaling pathway that are normally over-expressed in several cancer types (Jin and Song, 2014; Koehler and Schlupf, 2013). SOX9 overexpression has been identified to activate the Wnt/β-catenin signaling pathway thus promoting glioma metastasis (Liu and Liu, 2015). It has been revealed that over-expression of SOX9 in lung cancer cells could up-regulate the levels of p-Wnt and β-catenin (Guo and Xie, 2018). Similarly, miR-185 has been suggested to negatively target SOX9 thus leading to Wnt signaling inactivation and inhibition on NSCLC development (Lei, 2018). Besides, in addition to the inhibitory effect of DEX and miR-185 on OC cell growth *in vitro*, the *in vivo* studies presented a same trend. These results suggested that DEX treatment and over-expression of miR-185 could inhibit OC development with the involvement of SOX9 and the Wnt/β-catenin signaling pathway.

To sum up, the current study provided new insights that DEX could up-regulate miR-185 expression, which further inhibits SOX9 expression and then inactivate the Wnt/β-catenin signaling pathway, thus suppressing OC growth and development. Meanwhile, this finding provides evidence that DEX might serve as an anti-cancer agent more than just an anesthetic during cancer therapies. Hopefully, more studies would be applied in this field to validate our findings and to develop more novel therapies in treating OC.

## Acknowledgements

Not applicable.

## Funding

No funding was received.

## Availability of data and materials

All the data generated or analyzed during this study are included in this published article.

## Author contributions

HT is the guarantor of integrity of the entire study and contributed to the concepts and design of this study; LH contributed to the definition of intellectual content and literature research of this study; YMX contributed to the experimental studies and clinical studies; QJC contributed to the data analysis and statistical analysis; HT took charge of the manuscript preparation; LH and YMX contributed to the manuscript review. All authors read and approved the final manuscript.

## Ethical approval and consent to participate

The study was approved by the Clinical Ethical Committee of Guangzhou Medical University (Guangzhou, China). All experimental procedures were performed in line with the ethical guidelines for the study of experimental pain in conscious animals.

## Patient consent for publication

Not applicable.

## Competing interests

All authors declare that there is no conflict of interests in this study.

## Reference

Bareiss PM, Paczulla A, Wang H, Schairer R, Wiehr S, Kohlhofer U, Rothfuss OC, Fischer A, Perner S, Staebler A, Wallwiener D, Fend F, Fehm T, Pichler B, Kanz L, Quintanilla-Martinez L, Schulze-Osthoff K, Essmann F, Lengerke C. 2013. SOX2 expression associates with stem cell state in human ovarian carcinoma. Cancer Res 73, 5544–5555.

Basuli D, Tesfay L, Deng Z, Paul B, Yamamoto Y, Ning G, Xian W, McKeon F, Lynch M, Crum CP, Hegde P, Brewer M, Wang X, Miller LD, Dyment N, Torti FM, Torti SV. 2017. Iron addiction: a novel therapeutic target in ovarian cancer. Oncogene 36, 4089–4099.

Cai C, Wang H, He HH, Chen S, He L, Ma F, Mucci L, Wang Q, Fiore C, Sowalsky AG, Loda M, Liu XS, Brown M, Balk SP, Yuan X. 2013. ERG induces androgen receptor-mediated regulation of SOX9 in prostate cancer. J Clin Invest 123, 1109–1122.

Cai QH, Tang Y, Fan SH, Zhang ZF, Li H, Huang SQ, Wu DM, Lu J, Zheng YL. 2017. In vivo effects of dexmedetomidine on immune function and tumor growth in rats with ovarian cancer through inhibiting the p38MAPK/NF-kappaB signaling pathway. Biomed Pharmacother 95, 1830–1837.

Endesfelder S, Makki H, von Haefen C, Spies CD, Buhrer C, Sifringer M. 2017. Neuroprotective effects of dexmedetomidine against hyperoxia-induced injury in the developing rat brain. PLoS One 12, e0171498.

Freitas M, Alves V, Sarmento-Ribeiro AB, Mota-Pinto A. 2011. Combined effect of sodium selenite and docetaxel on PC3 metastatic prostate cancer cell line. Biochem Biophys Res Commun 408, 713–719.

Gao Y, Liu X, Li T, Wei L, Yang A, Lu Y, Zhang J, Li L, Wang S, Yin F. 2017. Cross-validation of genes potentially associated with overall survival and drug resistance in ovarian cancer. Oncol Rep 37, 3084–3092.

Guo YZ, Xie XL, Fu J, Xing GL. 2018. SOX9 regulated proliferation and apoptosis of human lung carcinoma cells by the Wnt/beta-catenin signaling pathway. Eur Rev Med Pharmacol Sci 22, 4898–4907.

Heimisdottir M. 2016. [Unlocking the full potential of Landspitali University Hospital - Icelandic healthcare at a crossroads. An outsiders view from McKinsey and Company[Editorial]]. Laeknabladid 102, 425.

Hua F, Li CH, Chen XG, Liu XP. 2018. Daidzein exerts anticancer activity towards SKOV3 human ovarian cancer cells by inducing apoptosis and cell cycle arrest, and inhibiting the Raf/MEK/ERK cascade. Int J Mol Med 41, 3485–3492.

Imam JS, Buddavarapu K, Lee-Chang JS, Ganapathy S, Camosy C, Chen Y, Rao MK. 2010. MicroRNA-185 suppresses tumor growth and progression by targeting the Six1 oncogene in human cancers. Oncogene 29, 4971–4979.

Ji J, Shi J, Budhu A, Yu Z, Forgues M, Roessler S, Ambs S, Chen Y, Meltzer PS, Croce CM, Qin LX, Man K, Lo CM, Lee J, Ng IO, Fan J, Tang ZY, Sun HC, Wang XW. 2009. MicroRNA expression, survival, and response to interferon in liver cancer. N Engl J Med 361, 1437–1447.

Jiang SS, Fang WT, Hou YH, Huang SF, Yen BL, Chang JL, Li SM, Liu HP, Liu YL, Huang CT, Li YW, Jang TH, Chan SH, Yang SJ, Hsiung CA, Wu CW, Wang LH, Chang IS. 2010. Upregulation of SOX9 in lung adenocarcinoma and its involvement in the regulation of cell growth and tumorigenicity. Clin Cancer Res 16, 4363–4373.

Jin XT, Song L, Zhao JY, Li ZY, Zhao MR, Liu WP. 2014. Dichlorodiphenyltrichloroethane exposure induces the growth of hepatocellular carcinoma via Wnt/beta-catenin pathway. Toxicol Lett 225, 158–166.

Koehler A, Schlupf J, Schneider M, Kraft B, Winter C, Kashef J. 2013. Loss of Xenopus cadherin-11 leads to increased Wnt/beta-catenin signaling and up-regulation of target genes c-myc and cyclin D1 in neural crest. Dev Biol 383, 132–145.

Lei Z, Shi H, Li W, Yu D, Shen F, Yu X, Lu D, Sun C, Liao K. 2018. miR185 inhibits nonsmall cell lung cancer cell proliferation and invasion through targeting of SOX9 and regulation of Wnt signaling. Mol Med Rep 17, 1742–1752.

Li F, Zhang L, Li W, Deng J, Zheng J, An M, Lu J, Zhou Y. 2015. Circular RNA ITCH has inhibitory effect on ESCC by suppressing the Wnt/beta-catenin pathway. Oncotarget 6, 6001–6013.

Li S, Ma Y, Hou X, Liu Y, Li K, Xu S, Wang J. 2015. MiR-185 acts as a tumor suppressor by targeting AKT1 in non-small cell lung cancer cells. Int J Clin Exp Pathol 8, 11854–11862.

Liu C, Li G, Ren S, Su Z, Wang Y, Tian Y, Liu Y, Qiu Y. 2017. miR-185-3p regulates the invasion and metastasis of nasopharyngeal carcinoma by targeting WNT2B in vitro. Oncol Lett 13, 2631–2636.

Liu H, Liu Z, Jiang B, Peng R, Ma Z, Lu J. 2015. SOX9 Overexpression Promotes Glioma Metastasis via Wnt/beta-Catenin Signaling. Cell Biochem Biophys 73, 205–212.

Liu M, Lang N, Chen X, Tang Q, Liu S, Huang J, Zheng Y, Bi F. 2011. miR-185 targets RhoA and Cdc42 expression and inhibits the proliferation potential of human colorectal cells. Cancer Lett 301, 151–160.

Lu B, Fang Y, Xu J, Wang L, Xu F, Xu E, Huang Q, Lai M. 2008. Analysis of SOX9 expression in colorectal cancer. Am J Clin Pathol 130, 897–904.

Lv J, Liang Y, Tu Y, Chen J, Xie Y. 2018. Hypoxic preconditioning reduces propofol-induced neuroapoptosis via regulation of Bcl-2 and Bax and downregulation of activated caspase-3 in the hippocampus of neonatal rats. Neurol Res 40, 767–773.

Ma F, Ye H, He HH, Gerrin SJ, Chen S, Tanenbaum BA, Cai C, Sowalsky AG, He L, Wang H, Balk SP, Yuan X. 2016. SOX9 drives WNT pathway activation in prostate cancer. J Clin Invest 126, 1745–1758.

Mohta M, Kalra B, Sethi AK, Kaur N. 2016. Efficacy of dexmedetomidine as an adjuvant in paravertebral block in breast cancer surgery. J Anesth 30, 252–260.

Nikpayam E, Tasharrofi B, Sarrafzadeh S, Ghafouri-Fard S. 2017. The Role of Long Non-Coding RNAs in Ovarian Cancer. Iran Biomed J 21, 3–15.

Ozga M, Aghajanian C, Myers-Virtue S, McDonnell G, Jhanwar S, Hichenberg S, Sulimanoff I. 2015. A systematic review of ovarian cancer and fear of recurrence. Palliat Support Care 13, 1771–1780.

Palanichamy JK, Rao DS. 2014. miRNA dysregulation in cancer: towards a mechanistic understanding. Front Genet 5, 54.

Palma E, Gonzalez V, Grunholz D, Landaeta M, Mallea M, Perez J, Armstrong T. 2017. [Shock as an adverse reaction to rituximab: Case report]. Rev Med Chil 145, 260–263.

Peng X, Chen K, Chen J, Fang J, Cui H, Zuo Z, Deng J, Chen Z, Geng Y, Lai W. 2016. Aflatoxin B1 affects apoptosis and expression of Bax, Bcl-2, and Caspase-3 in thymus and bursa of fabricius in broiler chickens. Environ Toxicol 31, 1113–1120.

Raspaglio G, Petrillo M, Martinelli E, Li Puma DD, Mariani M, De Donato M, Filippetti F, Mozzetti S, Prislei S, Zannoni GF, Scambia G, Ferlini C. 2014. Sox9 and Hif-2alpha regulate TUBB3 gene expression and affect ovarian cancer aggressiveness. Gene 542, 173–181.

Roberts SB, Wozencraft CP, Coyne PJ, Smith TJ. 2011. Dexmedetomidine as an adjuvant analgesic for intractable cancer pain. J Palliat Med 14, 371–373.

Ruan X, Liu A, Zhong M, Wei J, Zhang W, Rong Y, Liu W, Li M, Qing X, Chen G, Li R, Liao Y, Liu Q, Zhang X, Ren D, Wang Y. 2019. Silencing LGR6 Attenuates Stemness and Chemoresistance via Inhibiting Wnt/beta-Catenin Signaling in Ovarian Cancer. Mol Ther Oncolytics 14, 94–106.

Shanbehzadeh S, Salavati M, Talebian S, Khademi-Kalantari K, Tavahomi M. 2018. Attention demands of postural control in non-specific chronic low back pain subjects with low and high pain-related anxiety. Exp Brain Res 236, 1927–1938.

Thomsen MK, Ambroisine L, Wynn S, Cheah KS, Foster CS, Fisher G, Berney DM, Moller H, Reuter VE, Scardino P, Cuzick J, Ragavan N, Singh PB, Martin FL, Butler CM, Cooper CS, Swain A, Transatlantic Prostate G. 2010. SOX9 elevation in the prostate promotes proliferation and cooperates with PTEN loss to drive tumor formation. Cancer Res 70, 979–987.

Tian H, Hou L, Xiong Y, Cheng Q, Huang J. 2019. Effect of Dexmedetomidine-Mediated Insulin-Like Growth Factor 2 (IGF2) Signal Pathway on Immune Function and Invasion and Migration of Cancer Cells in Rats with Ovarian Cancer. Med Sci Monit 25, 4655–4664.

Varughese J, Cocco E, Bellone S, Bellone M, Todeschini P, Carrara L, Schwartz PE, Rutherford TJ, Pecorelli S, Santin AD. 2011. High-grade, chemotherapy-resistant primary ovarian carcinoma cell lines overexpress human trophoblast cell-surface marker (Trop-2) and are highly sensitive to immunotherapy with hRS7, a humanized monoclonal anti-Trop-2 antibody. Gynecol Oncol 122, 171–177.

Wang WX, Wu Q, Liang SS, Zhang XK, Hu Q, Chen QH, Huang HJ, Xu L, Lou FQ. 2018. Dexmedetomidine promotes the recovery of neurogenesis in aged mouse with postoperative cognitive dysfunction. Neurosci Lett 677, 110–116.

Wang X, Xu Y, Chen X, Xiao J. 2018. Dexmedetomidine Inhibits Osteosarcoma Cell Proliferation and Migration, and Promotes Apoptosis by Regulating miR-520a-3p. Oncol Res 26, 495–502.

Wang X, Zhao B, Li X. 2015. Dexmedetomidine attenuates isoflurane-induced cognitive impairment through antioxidant, anti-inflammatory and anti-apoptosis in aging rat. Int J Clin Exp Med 8, 17281–17288.

Weerink MAS, Struys M, Hannivoort LN, Barends CRM, Absalom AR, Colin P. 2017. Clinical Pharmacokinetics and Pharmacodynamics of Dexmedetomidine. Clin Pharmacokinet 56, 893–913.

Xiang Y, Ma N, Wang D, Zhang Y, Zhou J, Wu G, Zhao R, Huang H, Wang X, Qiao Y, Li F, Han D, Wang L, Zhang G, Gao X. 2014. MiR-152 and miR-185 co-contribute to ovarian cancer cells cisplatin sensitivity by targeting DNMT1 directly: a novel epigenetic therapy independent of decitabine. Oncogene 33, 378–386.

Xiao S, Li Y, Pan Q, Ye M, He S, Tian Q, Xue M. 2019. MiR-34c/SOX9 axis regulates the chemoresistance of ovarian cancer cell to cisplatin-based chemotherapy. J Cell Biochem 120, 2940–2953.

Xie L, Zhang Z, Tan Z, He R, Zeng X, Xie Y, Li S, Tang G, Tang H, He X. 2014. MicroRNA-124 inhibits proliferation and induces apoptosis by directly repressing EZH2 in gastric cancer. Mol Cell Biochem 392, 153–159.

Xu YB, D. QH, Zhang MY, Yun P, He CY. 2013. Propofol suppresses proliferation, invasion and angiogenesis by down-regulating ERK-VEGF/MMP-9 signaling in Eca-109 esophageal squamous cell carcinoma cells. Eur Rev Med Pharmacol Sci 17, 2486–2494.

Yam C, He Y, Zhang D, Chiam KH, Oliferenko S. 2011. Divergent strategies for controlling the nuclear membrane satisfy geometric constraints during nuclear division. Curr Biol 21, 1314–1319.

Zhang Y, Jia S, Gao T, Zhang R, Liu Z, Wang Y. 2018. Dexmedetomidine mitigate acute lung injury by inhibiting IL-17-induced inflammatory reaction. Immunobiology 223, 32–37.

